# The fallacy of global comparisons based on *per capita* measures

**DOI:** 10.1101/2023.05.19.541446

**Authors:** Lukáš Kratochvíl, Jan Havlíček

## Abstract

Social and environmental scientists often present comparisons across countries and policy makers use such comparisons to take evidence-based action. For a meaningful comparison among countries, one often needs to normalize the measure for differences in the population size. The first choice is usually to calculate *per capita* ratios. Such ratios, however, normalize the measure for differences in population size directly only under proportional increase of the measure with population size. Violation of this assumption frequently leads to misleading conclusions. To illustrate this issue, we compare *per capita* ratios with an approach based on regression, which eliminates many of the problems with ratios and allows for a straightforward data interpretation. The *per capita* measures in three global datasets (GDP, COVID-related mortality, and CO_2_ production) systematically overestimate values in countries with small populations, while countries with large populations tend to have misleadingly low *per capita* ratios due to the large denominators. Despite their biases, comparisons based on *per capita* ratios are still ubiquitous and are used for influential recommendations by various global institutions. Their continued use can cause significant damage when employed as evidence for policy actions and should be replaced by a more scientifically substantiated method, such as a regression-based approach.

## 1. Introduction

Social and environmental researchers often perform comparisons across countries (1,2) and policy makers use such comparisons to take evidence-based action (3,4). Absolute values are mostly not of much use because countries vary in their population size. For a meaningful comparison, one needs to control for differences in population size. To address this issue, the first choice is usually to calculate the ratio between the measure of interest and population size and then compare the values of *per capita* measures among countries. This intuitively appealing and seemingly straightforward procedure might, however, lead to misleading conclusions (5).

Some of the problems with ratios have been identified already at the dawn of statistics as a science (6,7). The major issue lies in the fact that the sole function yielding a constant when divided by its argument and thus removing the effect of a denominator is a linear function that goes through the origin of the coordinate system (figure 1). *Per capita* ratios can be directly used for comparisons across countries only when the measure of interest increases proportionally with population size. In all other cases, ratios ought to be interpreted with extreme caution due to their various nonintuitive statistical properties including: i) a frequently strong departure from normal distribution, ii) asymmetry with respect to changes in the numerator and denominator, and iii) scaling with the denominator (figure 1). Serious problems with ratios in comparative studies have been recognized in various branches of life sciences including morphometry (8), evolutionary genetics (9), and ecological stoichiometry (10), where the use of ratios has been largely abandoned and replaced by more appropriate procedures (7,11). Nevertheless, global comparisons of various aspects of economic performance, public health, or environmental impact are still frequently based on *per capita* indicators (12,13). Measures standardized in this way suffer from the abovementioned issues of ratios and frequently lead to worryingly misguided outcomes. Given the global impact of these measures, we believe there is an urgent need for alternatives. We propose an approach based on linear regression, which also provides simple and appealing measures but does not suffer from many of the problems which ratios have.

**Figure 1.**
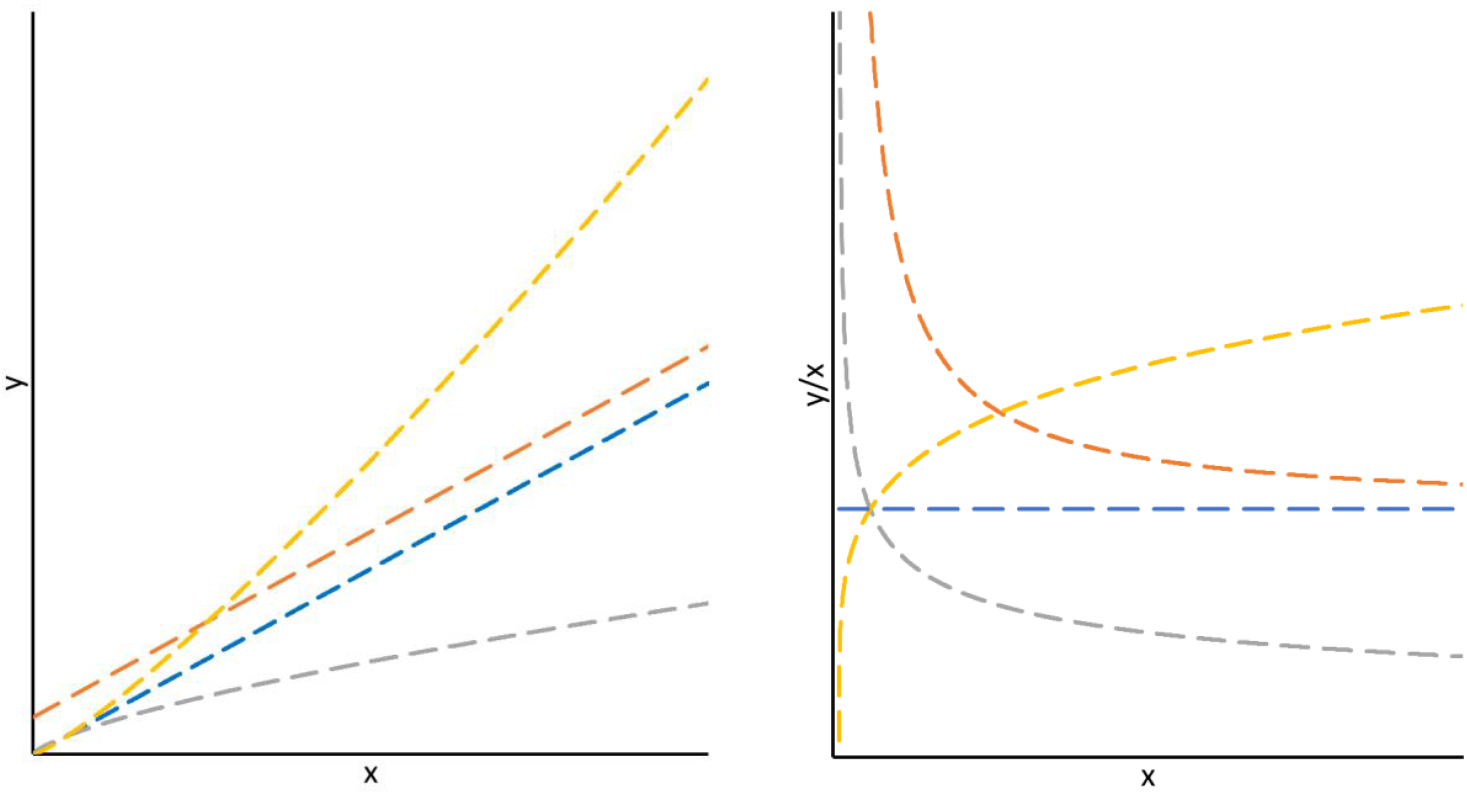
Exemplary increasing functions describing relationships between variables *x* (e.g., population size) and *y* (e.g., a country measure) (left) and dependence of the corresponding *y/x* ratios on the argument *x* (right). The only function giving a constant when divided by its argument is the linear function going through the origin (blue lines). Ratios thus properly standardize *y* for variability in *x* only when *y* increases in direct proportion to *x*. Note that in the other depicted cases, such as when the linear relationship between *y* and *x* does not go through the origin (orange) or an increase of *y* with x according to a power function (with power > 1 in yellow and between 0 and 1 in grey), the ratios have infinity (or minus infinity) limit as *x* tends to zero. As a consequence, even a small shift in *x* in small values produces a large change in the ratio *y/x*.

## 2. Three case studies

We compare the *per capita* ratios and the approach based on linear regression by applying them to three exemplar global comparisons in gross domestic product (GDP; data from https://datacatalog.worldbank.org), COVID-19 related mortality (data from https://www.worldometers.info/coronavirus/#countries), and CO_2_ production (data from https://www.worldometers.info/co2-emissions/). We selected these three variables because they are widely used and affect political decisions in economy, health systems, environment conservation, as well as other areas (13,14). Specialists in the relevant fields extensively discuss the pros and cons of these measures. For instance, it is largely recognized that GDP need not be the best proxy for country’s economic performance and progress (15,16). Nevertheless, it was shown that GDP *per capita* is closely correlated with the more sophisticated measures such as Sustainable Development Goals index, Human Development Index and Environmental Performance Index (17). Similarly, data on COVID-19-related mortality may suffer from systematic inaccuracies related to differences in data gathering and publication among countries (18,19). At present, we shall focus on the problems of *per capita* comparisons and leave aside other questions concerning the suitability of particular measures (for a discussion see (16,20)).

The *per capita* expressions in all three measures (GDP, COVID-19 mortality, CO_2_ production) show a pattern typical of many ratios. Firstly, the ratios substantially deviate from normal distribution (electronic supplementary material figure S1). The mean *per capita* value, while often used to describe the central tendency, is therefore not an informative representative of the dataset. Secondly, the ratios exhibit enormous variability among countries with small populations; this variability then sharply decreases with countries’ increasing population size (figure 2). Small changes in population size thus have disproportionately larger effects in countries with small populations. Thirdly, the GDP and the COVID-19 related mortality data show a strong negative correlation with population size (Spearman’s ρ = -0.32, p < 0.00001, and ρ = -0.20, p < 0.004, respectively). The data on GDP and COVID-19 related mortality thus do not increase proportionally with population size and the *per capita* ratios do not adequately standardize for differences in population size. This is further reflected in a strong correlation between country ranking according to *per capita* GDP and *per capita* COVID-19 deaths rates on the one hand and population size on the other hand (Spearman’s ρ = 0.32, p = 0.000002, and ρ = 0.20, p = 0.004, respectively, figure 3). In contrast, the correlation between *per capita* CO_2_ emissions and population size is not statistically significant (ρ = -0.11, p = 0.10). Consequently, the correlation between a ranking based on *per capita* CO_2_ emissions and log-transformed population size is not statistically significant (ρ = 0.09, p = 0.21). Nevertheless, the shifts in *per capita* ratios in CO_2_ emissions demonstrate a strong asymmetry with shifts in population size (electronic supplementary material figure S1), suggesting that the *per capita* ratio is not an optimal index for this measure either.

**Figure 2.**
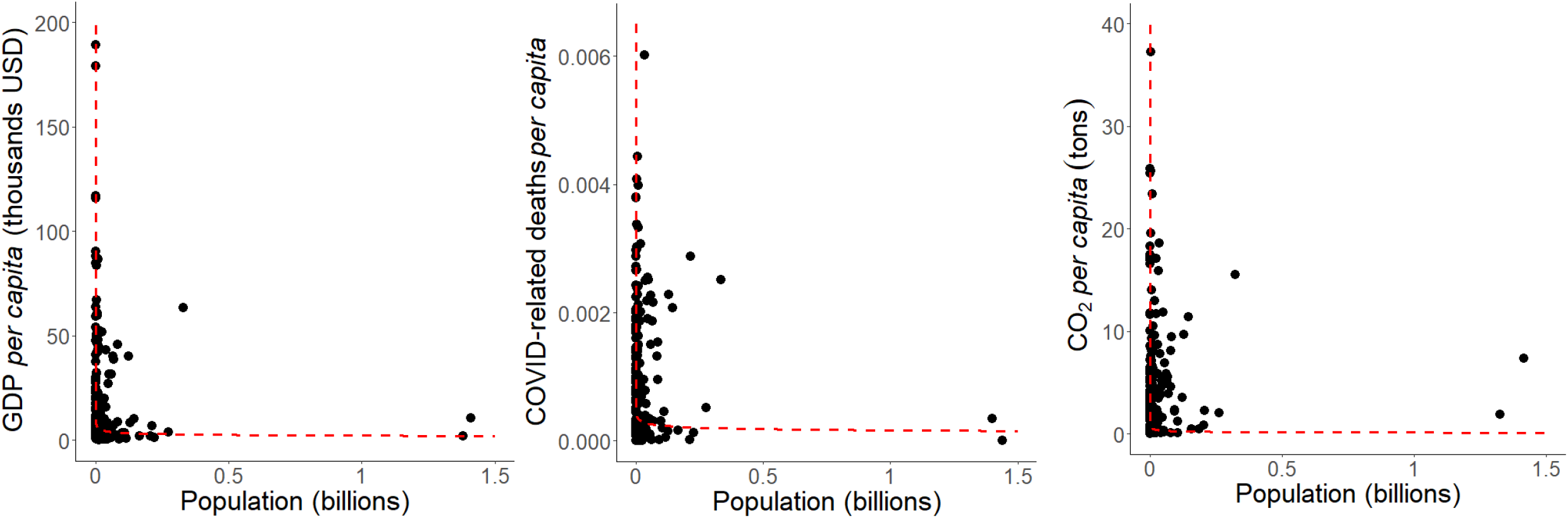
Plot of *per capita* measures against population size across countries. The *per capita* ratios in all three measures (GDP, COVID-related deaths, and CO_2_ emissions) exhibit enormous variability among countries with small populations. This variability then decreases with increasing country population size. This is due to the infinity limits of the underlying functions as population size approaches zero. The red lines represent predictions for ratios derived from the back-transformation of the linear relationship between logarithmically transformed values of each measure and populations size depicted in the Figure 3b.

**Figure 3.**
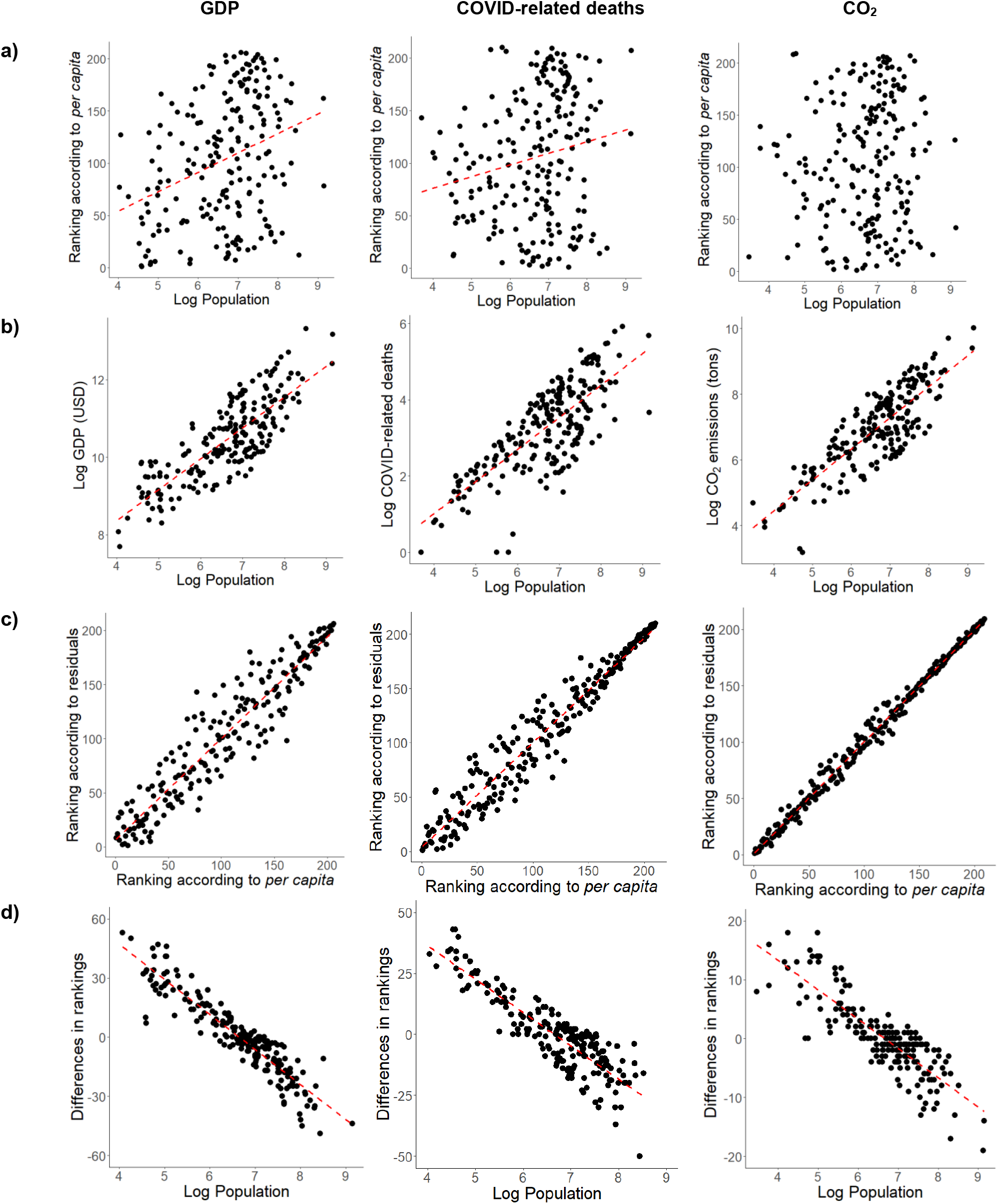
Scatterplots comparing country rankings based on *per capita* ratios and residuals. a) Rankings according to *per capita* ratios significantly increase with population size in two out of three measures, demonstrating that the ratios do not really control for population size. b) The increase of measures with population size. **c)** Rankings based on ratios and residuals are correlated. **d)** Differences between rankings based on *per capita* ratios and residuals highly correlate with population size, which indicates that *per capita* ratios systematically overestimate values in countries with small populations and underestimate them in countries with large populations.

The assumption of proportionality of a measure to the population size is rarely tested and even less frequently met. Researchers who use the *per capita* ratio to normalize a measure to population size should be aware that a violation of the proportionality assumption leads to severe biases when *per capita* ratios are used for direct comparisons among countries. A correct approach to global comparisons should not be based on unrealistic and often violated assumptions: it should consider the general trend of the data. We cannot stress enough that before using any analytical technique, one should carefully check the trends in a given dataset including its distribution, central tendency, and effect of outliers. The properties of data one ought to consider carefully are well-known and include non-linearity, homoscedasticity, and leverage (7).

In all three examples analyzed here, the linear function of log_10_-transformed measure to log_10_-transformed population size describes the general trend well. These functions explain 65%, 59%, and 68% of total variance, demonstrating that total GDP, COVID-related mortality, and CO_2_ production, respectively, scale with population size, and that a standardization for population size is necessary for comparisons. The addition of a quadratic term to each of these relationships did not significantly improve the model’s explanatory power (for details, see electronic supplementary material). We thus further neglect a slight nonlinearity in the log-log relationships.

Importantly, these linear approximations on the log-log scale explain the behaviour of the *per capita* ratios very well. To illustrate it, we back-transformed the linear functions of logarithmically transformed values and depicted the ratios of values predicted by the back-transformed power functions for a given population size (figure 2). The predicted *per capita* measures approach infinity as the population size approaches zero, which is responsible for the strong asymmetry in *per capita* ratios. Most notably, due to the steepness of the functions as they approach zero, a small shift in the population size or the measure of interest in countries with a small population leads to an enormous shift in the *per capita* ratio. In contrast, even a large change in population size or the measure of interest in countries with large populations has little effect on the *per capita* ratios and their ratios are moreover in general relatively low. Nevertheless, regardless of the precise underlying function, in all cases the *per capita* ratios are clearly inappropriate direct standardization for differences in population size among countries.

## 3. Differences in country ranking between the two approaches

Instead of the highly problematic *per capita* ratios for comparison among countries, one may employ residuals from the log-log linear regression. These residuals have a straightforward interpretation: their values +1.0 and -1.0 correspond to ten times higher and ten times lower values of a measure in comparison to the value expected for a country of a particular population size (residual 0). The similarities and differences between the *per capita* approach and the regression approach can be exemplified by comparing the rankings of individual countries in a particular measure. Rankings based on the two approaches were in all three measures highly correlated, which reflects the enormous differences among countries (figure 3), but we also found some important differences between the rankings. None of the rankings based on the linear regression approach correlated with the population size, which is exactly what one would expect from a procedure standardizing for differences in population size. The direction of shifts in the rankings in all three measures highly correlated with country population sizes (figure 3), which is what one would expect from the relationship between *per capita* ratio and population size (figure 2). In particular it can be expected that some small countries will have misleadingly large ratios because of the small denominator (population size), while large countries will have misleadingly low ratios due to large denominators substantially decreasing their metrics. We recorded the largest change in the GDP ranking relative to population size in Nauru, which moved 66 positions down, while India moved 64 positions up in the ranking based on the regression approach as compared to the *per capita* measures (electronic supplementary material table S1). In fact, all 29 countries with population above 50 million moved up (on average 29 positions, which is considerable). The effect of large denominators on the *per capita* ratio can be exemplified in China and India, the two most populous countries in the world. According to the GDP *per capita* measure, both are within a broad middle of the ranking: China ranks 78^th^ and India even 162^nd^ among 206 countries. But when we compare countries without the bias introduced by the behaviour of ratios, India (with residual -0.04) reaches 91% of GDP expected for a country of its population size, and China (residual 0.70) has an over five times higher GDP than predicted for a country of its population size. In ranking according to residuals, India is around the middle of the ranking (98^th^ place), while China is among the 17% of highest-scoring countries (34^th^ place). Also, we can notice that only small population countries are represented in the top ten of the GDP ranking according to *per capita* ratios, while in ranking according to residuals, the top ten countries are much more balanced with respect to their population size (figure 4).

**Figure 4.**
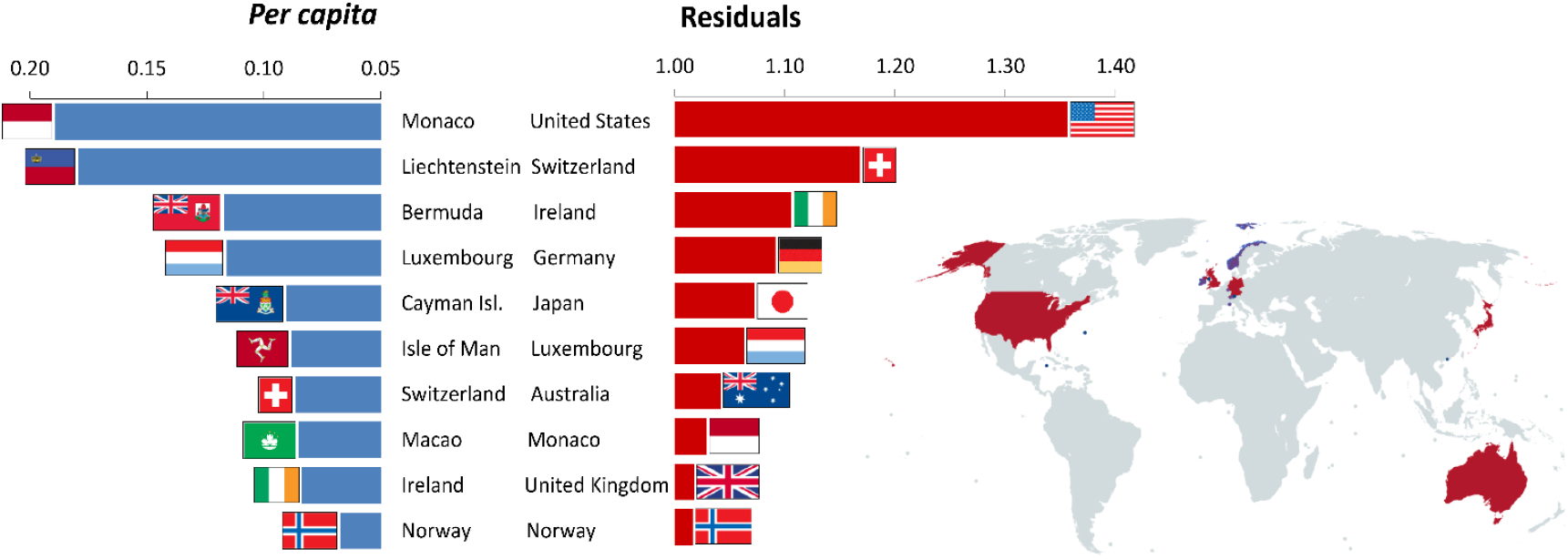
Top 10 countries in GDP controlled for population size highly differ in rankings according to *per capita* measures and residuals. Note that only countries with small population are represented among the top ten according to the *per capita* ratios (given as USD *per capita*), which reflects an overestimation of their measures by this procedure.

Similarly as for the GDP, three countries of small population size (Gibraltar, San Marino, and Liechtenstein) versus two countries with large population (Indonesia and India) showed the largest change in COVID-related mortality rankings, with the small countries moving up (in terms of higher mortality) and the large ones down (electronic supplementary material table S2). Two countries with a large population (USA and Brazil), where the situation with COVID-related mortality was apparently dramatic (21,22), took in the regression-based ranking the second and the third position (in highest morality) right after Peru, where COVID-related mortality was so disastrous (23) that it headed the rankings based on both approaches. Again, the much lower positions of USA and Brazil in the *per capita* ranking should be attributed to their large population size, that is, the large denominator in the calculation of the *per capita* ratio.

In CO_2_ emission ranking, the situation was analogous but the changes in ranking were smaller. Still, the most notable movements between the rankings in the opposite directions made Palau and Seychelles, which moved down, versus India and Brazil, which moved up (electronic supplementary material table S3). Countries with population over 50 million inhabitants moved on average 7 positions up, meaning their production of CO_2_ emissions standardized for population size is in a global comparison comparatively much higher than one would assume based on the *per capita* ranking.

## 4. Conclusions

We are not the first to draw attention to the biases introduced by the use of ratios and the need for alternatives. For instance, Ranta et al. (11) claimed that a regression-based approach amounts to reinventing the wheel in their instructive demonstration of problems with ratios in comparisons of sexual size dimorphism among animals. Still, the use of *per capita* measures, an exemplary case of ratios, remains ubiquitous. For instance, the World Bank classifies the world’s economies in four income groups – low, lower-middle, upper-middle, and high-income countries – based on *per capita* gross national income. Therefore, we feel compelled to stress that a continued use of *per capita* measures is not only of a limited informative value but can result in misleading conclusions and cause extensive damage when employed as evidence for policy actions. The alternative offered here, the regression-based approaches, are computationally simple and can be flexibly adjusted for general trends in data. The interpretation of such comparisons is straightforward, and the results can be thus easily communicated to the general public. We believe that policy makers, practitioners, and influential organizations, such as the Food and Agriculture Organization (24), the World Health Organization (25), World Bank (14), International Monetary Fund (26), Eurostat (27), and the Organisation for Economic Co-operation and Development (28) as well as various widely used information sources such as Wikipedia (29), could adopt it easily and abandon the *per capita* approach, which appears to standardize well for a population size but in fact provides distorted insights.

## Authors’ contributions

Both authors contributed to the conception of the study, manuscript preparation and its final revision. LK performed the data analysis.

## Acknowledgements

We would like to express our gratitude to Barbora Straková, Jasna Vukić and Petr Tureček for comments on previous versions of the manuscript, and Anna Pilátová for English proofreading.

## References

1. Ravallion M. 2020 On measuring global poverty. Annu. Rev. Econom. 12, 167–188. (doi:10.1146/annurev-economics-081919-022924)

2. Black RE, Morris SS, Bryce J. 2003 Where and why are 10 million children dying every year? Lancet 361, 2026–2034. (doi:10.1016/S0140-6736(03)13779-8)

3. Anell A, Willis M. 2000 International comparison of health care systems using resource profiles. Bull. World Health Organ. 78, 770–778.

4. Ridley M, Rao G, Schilbach F, Patel V. 2020 Poverty, depression, and anxiety: Causal evidence and mechanisms. Science 370, eaay0214. (doi:10.1126/science.aay0214)

5. Silva WTAF. 2020 Per capita death and infection rates should be avoided in international comparisons. Public Health 186, 18–19. (doi:10.1016/j.puhe.2020.06.038)

6. Pearson K. 1897 Mathematical contributions to the theory of evolution.—on a form of spurious correlation which may arise when indices are used in the measurement of organs. Proc. R. Soc. London 60, 489–498.

7. Sokal R, Rohlf F. 1969 Biometry: the principles and practices of statistics in biological research. San Fransisco: Freeman.

8. Lolli L, Batterham AM, Kratochvíl L, Flegr J, Weston KL, Atkinson G. 2017 A comprehensive allometric analysis of 2^nd^ digit length to 4^th^ digit length in humans. Proc. R. Soc. B Biol. Sci. 284, 20170356. (doi:10.1098/rspb.2017.0356)

9. Kratochvíl L, Rovatsos M. 2022 Ratios of non-synonymous to synonymous mutations can be misleading for detecting selection. Curr. Biol. 32, R28–R30.

10. Isles PDF. 2020 The misuse of ratios in ecological stoichiometry. Ecology 101, 1–7.

11. Ranta E, Laurila A, Elmberg J. 1994 Reinventing the wheel: analysis of sexual dimorphism in body size. Oikos 70, 313–321.

12. Li X, Lin B. 2013 Global convergence in per capita CO2 emissions. Renew. Sustain. Energy. Rev. 24, 357–363. (doi:10.1016/j.rser.2013.03.048)

13. WHO. 2019 Global Spending on Health: A World in Transition 2019. Global Report. 68 p. https://www.who.int/health_financing/documents/health-expenditure-report-2019.pdf?ua=1

14. World Bank. World Development Report 2022: Finance for an equitable recovery. Washington, D.C.: World Bank Publications; 2022.

15. Stiglitz JE, Sen A, Fitoussi J-P. 2009 Report by the Commission on the Measurement of Economic Performance and Social Progress.

16. Costanza R, Hart M, Posner S, Talberth J. 2009 Beyond GDP : The Need for New Measures of Progress Beyond GDP : The Need for New Measures of Progress. Bost Univ. 4, 1–47.

17. Schmidt-Traub G, Kroll C, Teksoz K, Durand-Delacre D, Sachs JD. 2017 National baselines for the sustainable development goals assessed in the SDG Index and dashboards. Nat. Geosci. 10, 547–555. (doi:10.1038/ngeo2985)

18. Corrao G, Rea F, Blangiardo GC. 2021 Lessons from COVID-19 mortality data across countries. J. Hypertens. 398, 56–60. (doi:10.1097/HJH.0000000000002833)

19. Middelburg RA, Rosendaal FR. 2020 COVID-19: How to make between-country comparisons. Int. J. Infect. Dis. 96, 477–481. (doi:10.1016/j.ijid.2020.05.066)

20. Giannetti BF, Agostinho F, Almeida CMVB, Huisingh D. 2015 A review of limitations of GDP and alternative indices to monitor human wellbeing and to manage eco-system functionality. J. Clean. Prod. 87, 11–25. (doi:10.1016/j.jclepro.2014.10.051)

21. Ranzani OT, Bastos LSL, Gelli JGM, Marchesi JF, Baião F, Hamacher S, et al. 2021 Characterisation of the first 250 000 hospital admissions for COVID-19 in Brazil: a retrospective analysis of nationwide data. Lancet Respir. Med. 9, 407–418. (doi:10.1016/S2213-2600(20)30560-9)

22. Jin J, Agarwala N, Kundu P, Harvey B, Zhang Y, Wallace E, et al. 2021 Individual and community-level risk for COVID-19 mortality in the United States. Nat. Med. 27, 264–269. (doi:10.1038/s41591-020-01191-8)

23. Sempé L, Lloyd-Sherlock P, Martínez R, Ebrahim S, McKee M, Acosta E. 2021 Estimation of all-cause excess mortality by age-specific mortality patterns for countries with incomplete vital statistics: a population-based study of the case of Peru during the first wave of the COVID-19 pandemic. Lancet Reg. Health Am. 2, 100039. (doi:10.1016/j.lana.2021.100039)

24. Food and Agriculture Organization. https://www.fao.org/faostat/

25. WHO COVID 19. https://covid19.who.int/

26. International Monetary Fund. https://www.imf.org/external/datamapper/NGDP_RPCH@WEO/OEMDC/ADVEC/WEOWORLD

27. Eurostat. https://ec.europa.eu/eurostat/databrowser/view/sdg_08_10/default/table?lang=en

28. The Organisation for Economic Co-operation and Development. https://stats.oecd.org/

29. Wikipedia. https://en.wikipedia.org/wiki/Gross_domestic_product

