## Supplementary material for "The fallacy of global comparisons based on *per capita* measures"

**Electronic supplementary material**

***Test of non-linearity in the log-log relationships***

We compared the explanatory power of the linear and non-linear models between log-transformed values of the measures and log-transformed population size using Akaike information criterion (AIC) values. When ΔAIC was <2, the models were considered equivalent, but priority was given to the simplest model while a more complex model would be considered supported when ΔAIC > 2. To test non-linearity, we compared the linear model and the model with the quadratic term added (the best model in bold):

**Model	df	AIC**

Log(GDP) ~ Log(Population)	**3	382.04**

Log(GDP) ~ Log(Population) + Log(Population)^2	4	381.35

Log(COVID-related deaths) ~ Log(Population)	**3	476.92**

Log(COVID-related deaths) ~ Log(Population) + Log(Population)^2	4	478.87

Log(CO_2_ emissions) ~ Log(Population)	**3	427.95**

Log(CO_2_ emissions) ~ Log(Population) + Log(Population)^2	4	429.61

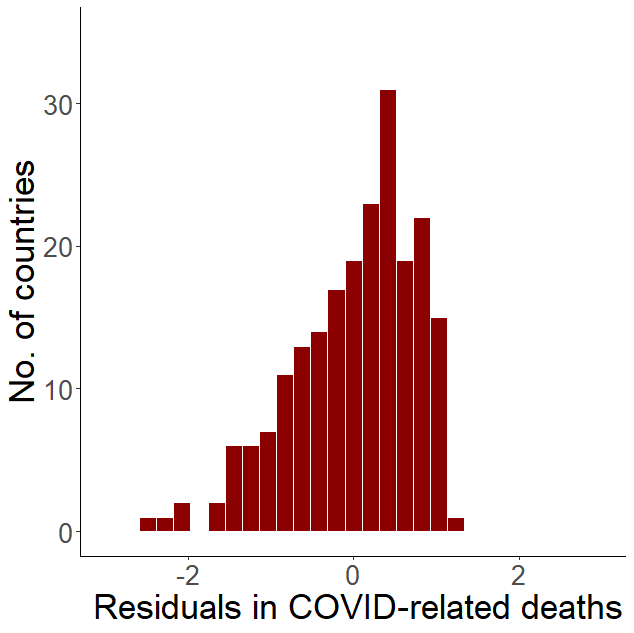

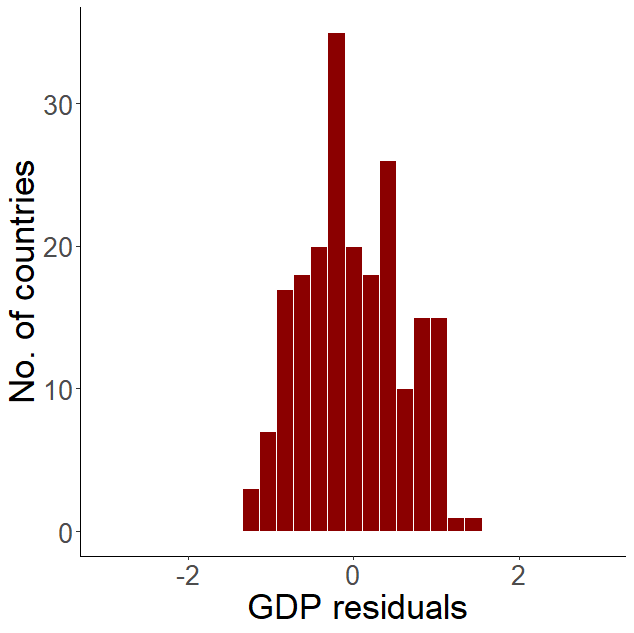

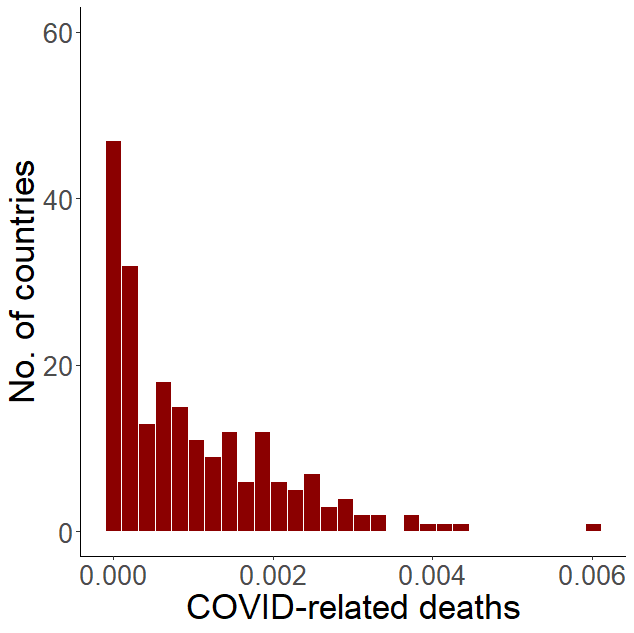

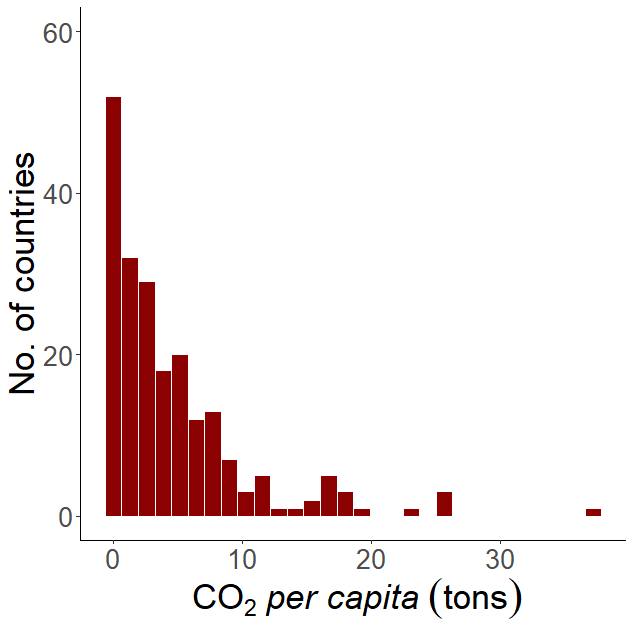

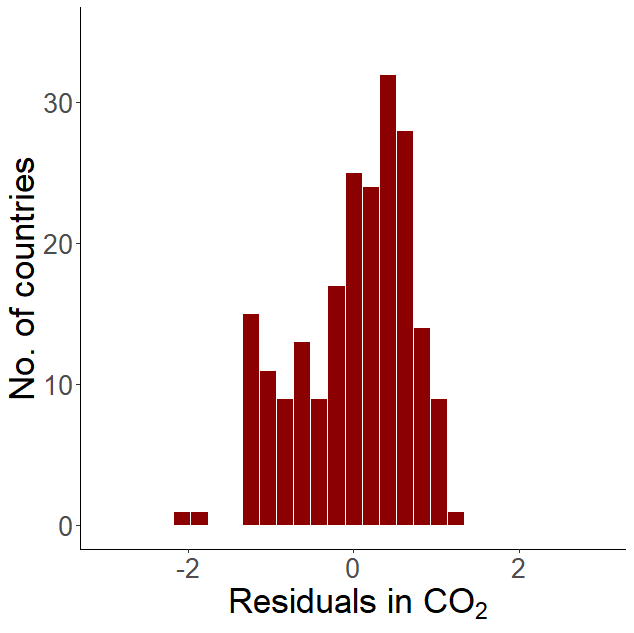

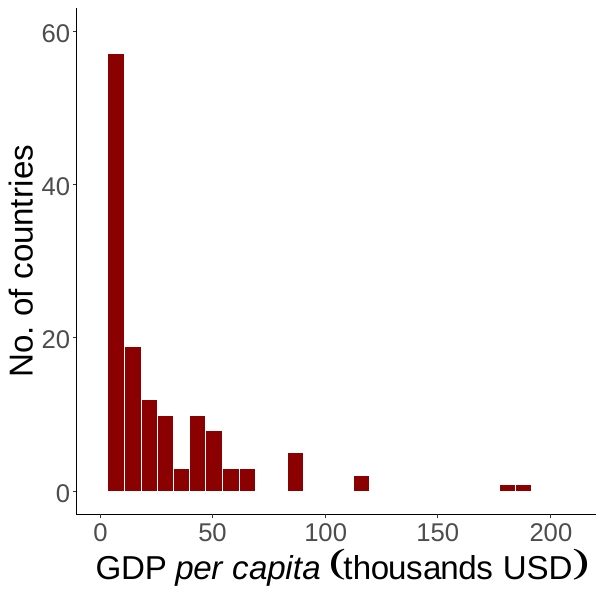

**Figure S1.** Histograms of *per capita* ratios and residuals from population size in GDP, COVID-related deaths, and CO_2_ emissions.

**Table S1.** A list of countries ranked according to their GDP standardized by application of linear regression, the use of *per capita* ratios, and the difference between the two approaches.

| **Country** | **Ranking based on residuals** | **Ranking based on *per capita* ratios** | **Difference in order** |
| --- | --- | --- | --- |
| United States | 1 | 12 | -11 |
| Switzerland | 2 | 7 | -5 |
| Ireland | 3 | 9 | -6 |
| Germany | 4 | 25 | -21 |
| Japan | 5 | 33 | -28 |
| Luxembourg | 6 | 4 | +2 |
| Australia | 7 | 19 | -12 |
| Monaco | 8 | 1 | +7 |
| United Kingdom | 9 | 32 | -23 |
| Norway | 10 | 10 | 0 |
| The Netherlands | 11 | 17 | -6 |
| Liechtenstein | 12 | 2 | +10 |
| France | 13 | 34 | -21 |
| Canada | 14 | 28 | -14 |
| Denmark | 15 | 13 | +2 |
| Singapore | 16 | 14 | +2 |
| Sweden | 17 | 18 | -1 |
| Macao SAR, China | 18 | 8 | +10 |
| Austria | 19 | 22 | -3 |
| Belgium | 20 | 26 | -6 |
| Italy | 21 | 39 | -18 |
| Korea, Rep. | 22 | 40 | -18 |
| Hong Kong SAR, China | 23 | 24 | -1 |
| Finland | 24 | 21 | +3 |
| Israel | 25 | 27 | -2 |
| United Arab Emirates | 26 | 29 | -3 |
| Bermuda | 27 | 3 | +24 |
| Qatar | 28 | 20 | +8 |
| Spain | 29 | 46 | -17 |
| New Zealand | 30 | 30 | 0 |
| Isle of Man | 31 | 6 | +25 |
| Cayman Islands | 32 | 5 | +27 |
| Iceland | 33 | 15 | +18 |
| China | 34 | 78 | -44 |
| Kuwait | 35 | 38 | -3 |
| Saudi Arabia | 36 | 54 | -18 |
| Puerto Rico | 37 | 37 | 0 |
| Czech Republic | 38 | 50 | -12 |
| Portugal | 39 | 52 | -13 |
| Faroe Islands | 40 | 11 | +29 |
| Poland | 41 | 63 | -22 |
| Greenland | 42 | 16 | +26 |
| Slovenia | 43 | 47 | -4 |
| Greece | 44 | 59 | -15 |
| Russian Federation | 45 | 80 | -35 |
| Slovak Republic | 46 | 58 | -12 |
| Guam | 47 | 36 | +11 |
| Hungary | 48 | 62 | -14 |
| Bahrain | 49 | 51 | -2 |
| Estonia | 50 | 49 | +1 |
| Lithuania | 51 | 56 | -5 |
| Malta | 52 | 44 | +8 |
| Andorra | 53 | 31 | +22 |
| Chile | 54 | 71 | -17 |
| San Marino | 55 | 23 | +32 |
| Virgin Islands (U.S.) | 56 | 35 | +21 |
| The Bahamas | 57 | 43 | +14 |
| Romania | 58 | 72 | -14 |
| Brunei Darussalam | 59 | 45 | +14 |
| Mexico | 60 | 89 | -29 |
| Malaysia | 61 | 79 | -18 |
| Turkey | 62 | 87 | -25 |
| Oman | 63 | 67 | -4 |
| Cyprus | 64 | 57 | +7 |
| Brazil | 65 | 100 | -35 |
| Latvia | 66 | 60 | +6 |
| Uruguay | 67 | 64 | +3 |
| Aruba | 68 | 41 | +27 |
| Croatia | 69 | 70 | -1 |
| Argentina | 70 | 88 | -18 |
| Thailand | 71 | 96 | -25 |
| Costa Rica | 72 | 74 | -2 |
| Panama | 73 | 73 | 0 |
| Trinidad and Tobago | 74 | 65 | +9 |
| Kazakhstan | 75 | 85 | -10 |
| Sint Maarten (Dutch part) | 76 | 42 | +34 |
| Cuba | 77 | 84 | -7 |
| Bulgaria | 78 | 81 | -3 |
| Curacao | 79 | 55 | +24 |
| Peru | 80 | 105 | -25 |
| Turks and Caicos Islands | 81 | 48 | +33 |
| Indonesia | 82 | 131 | -49 |
| Colombia | 83 | 109 | -26 |
| Dominican Republic | 84 | 95 | -11 |
| Barbados | 85 | 66 | +19 |
| South Africa | 86 | 111 | -25 |
| Northern Mariana Islands | 87 | 53 | +34 |
| Serbia | 88 | 91 | -3 |
| Turkmenistan | 89 | 92 | -3 |
| Belarus | 90 | 104 | -14 |
| Ecuador | 91 | 108 | -17 |
| St. Kitts and Nevis | 92 | 61 | +31 |
| Egypt, Arab Rep. | 93 | 135 | -42 |
| Mauritius | 94 | 86 | +8 |
| Antigua and Barbuda | 95 | 69 | +26 |
| Iraq | 96 | 126 | -30 |
| Philippines | 97 | 142 | -45 |
| India | 98 | 162 | -64 |
| Gabon | 99 | 98 | +1 |
| Guatemala | 100 | 117 | -17 |
| Botswana | 101 | 101 | 0 |
| Bosnia and Herzegovina | 102 | 106 | -4 |
| Equatorial Guinea | 103 | 97 | +6 |
| Ukraine | 104 | 137 | -33 |
| Paraguay | 105 | 112 | -7 |
| Lebanon | 106 | 113 | -7 |
| Algeria | 107 | 141 | -34 |
| Seychelles | 108 | 76 | +32 |
| Vietnam | 109 | 148 | -39 |
| Montenegro | 110 | 90 | +20 |
| Sri Lanka | 111 | 134 | -23 |
| Jordan | 112 | 121 | -9 |
| North Macedonia | 113 | 107 | +6 |
| Azerbaijan | 114 | 124 | -10 |
| Guyana | 115 | 99 | +16 |
| Maldives | 116 | 93 | +23 |
| St. Lucia | 117 | 83 | +34 |
| Palau | 118 | 68 | +50 |
| Albania | 119 | 110 | +9 |
| American Samoa | 120 | 75 | +45 |
| Morocco | 121 | 147 | -26 |
| Nigeria | 122 | 158 | -36 |
| Grenada | 123 | 82 | +41 |
| Suriname | 124 | 103 | +21 |
| Jamaica | 125 | 116 | +9 |
| Iran, Islamic Rep. | 126 | 155 | -29 |
| Georgia | 127 | 122 | +5 |
| Bangladesh | 128 | 160 | -32 |
| El Salvador | 129 | 132 | -3 |
| Moldova | 130 | 118 | +12 |
| Tunisia | 131 | 140 | -9 |
| Libya | 132 | 133 | -1 |
| Armenia | 133 | 123 | +10 |
| Bolivia | 134 | 144 | -10 |
| Mongolia | 135 | 130 | +5 |
| Namibia | 136 | 125 | +11 |
| Kosovo | 137 | 120 | +17 |
| Fiji | 138 | 114 | +24 |
| Ghana | 139 | 153 | -14 |
| St. Vincent and the Grenadines | 140 | 94 | +46 |
| Cote d'Ivoire | 141 | 154 | -13 |
| West Bank and Gaza | 142 | 143 | -1 |
| Nauru | 143 | 77 | +66 |
| Kenya | 144 | 164 | -20 |
| Papua New Guinea | 145 | 150 | -5 |
| Lao PDR | 146 | 151 | -5 |
| Angola | 147 | 163 | -16 |
| Honduras | 148 | 152 | -4 |
| Dominica | 149 | 102 | +47 |
| Belize | 150 | 119 | +31 |
| Pakistan | 151 | 176 | -25 |
| Eswatini | 152 | 139 | +13 |
| Uzbekistan | 153 | 165 | -12 |
| Djibouti | 154 | 138 | +16 |
| Myanmar | 155 | 172 | -17 |
| Tonga | 156 | 115 | +41 |
| Bhutan | 157 | 145 | +12 |
| Cameroon | 158 | 169 | -11 |
| Samoa | 159 | 128 | +31 |
| Nicaragua | 160 | 161 | -1 |
| Congo, Rep. | 161 | 159 | +2 |
| Cabo Verde | 162 | 146 | +16 |
| Cambodia | 163 | 168 | -5 |
| Senegal | 164 | 170 | -6 |
| Ethiopia | 165 | 183 | -18 |
| Tanzania | 166 | 182 | -16 |
| Nepal | 167 | 179 | -12 |
| Mauritania | 168 | 167 | +1 |
| Micronesia, Fed. Sts. | 169 | 136 | +33 |
| Marshall Islands | 170 | 129 | +41 |
| Vanuatu | 171 | 149 | +22 |
| Benin | 172 | 174 | -2 |
| Solomon Islands | 173 | 156 | +17 |
| Guinea | 174 | 175 | -1 |
| Zimbabwe | 175 | 180 | -5 |
| Haiti | 176 | 177 | -1 |
| Zambia | 177 | 181 | -4 |
| Uganda | 178 | 189 | -11 |
| Kyrgyz Republic | 179 | 178 | +1 |
| Tuvalu | 180 | 127 | +53 |
| Mali | 181 | 187 | -6 |
| Sao Tome and Principe | 182 | 157 | +25 |
| Yemen, Rep. | 183 | 191 | -8 |
| Burkina Faso | 184 | 188 | -4 |
| Timor-Leste | 185 | 173 | +12 |
| Togo | 186 | 184 | +2 |
| Congo, Dem. Rep. | 187 | 199 | -12 |
| Comoros | 188 | 171 | +17 |
| Tajikistan | 189 | 186 | +3 |
| Rwanda | 190 | 190 | 0 |
| Sudan | 191 | 196 | -5 |
| Malawi | 192 | 194 | -2 |
| Chad | 193 | 195 | -2 |
| Kiribati | 194 | 166 | +28 |
| Niger | 195 | 198 | -3 |
| Afghanistan | 196 | 200 | -4 |
| Lesotho | 197 | 185 | +12 |
| Madagascar | 198 | 201 | -3 |
| The Gambia | 199 | 192 | +7 |
| Mozambique | 200 | 204 | -4 |
| Guinea-Bissau | 201 | 193 | +8 |
| Liberia | 202 | 197 | +5 |
| Sierra Leone | 203 | 202 | +1 |
| Central African Republic | 204 | 203 | +1 |
| Somalia | 205 | 205 | 0 |
| Burundi | 206 | 206 | 0 |

**Table S2.** A list of countries ranked according to their COVID-19 related mortality standardized by the use of linear regression, by *per capita* ratios, and the difference between the two approaches.

| **Country Name** | **Ranking based on residuals** | **Ranking based on *per capita* ratios** | **Difference in order** |
| --- | --- | --- | --- |
| Peru | 1 | 1 | 0 |
| Brazil | 2 | 14 | -12 |
| USA | 3 | 19 | -16 |
| Bulgaria | 4 | 2 | +2 |
| Hungary | 5 | 4 | +1 |
| Mexico | 6 | 26 | -20 |
| Bosnia and Herzegovina | 7 | 3 | +4 |
| Romania | 8 | 9 | -1 |
| Czech Republic | 9 | 8 | +1 |
| Colombia | 10 | 20 | -10 |
| Argentina | 11 | 18 | -7 |
| Russia | 12 | 32 | -20 |
| Poland | 13 | 21 | -8 |
| Italy | 14 | 27 | -13 |
| North Macedonia | 15 | 6 | +9 |
| United Kingdom | 16 | 30 | -14 |
| Georgia | 17 | 7 | +10 |
| Ukraine | 18 | 29 | -11 |
| Slovakia | 19 | 11 | +8 |
| Croatia | 20 | 10 | +10 |
| France | 21 | 40 | -19 |
| Belgium | 22 | 23 | -1 |
| Montenegro | 23 | 5 | +18 |
| Spain | 24 | 38 | -14 |
| Chile | 25 | 34 | -9 |
| Lithuania | 26 | 15 | +11 |
| Armenia | 27 | 16 | +11 |
| Tunisia | 28 | 31 | -3 |
| Paraguay | 29 | 25 | +4 |
| Iran | 30 | 54 | -24 |
| Slovenia | 31 | 17 | +14 |
| Moldova | 32 | 24 | +8 |
| Ecuador | 33 | 41 | -8 |
| South Africa | 34 | 55 | -21 |
| Greece | 35 | 36 | -1 |
| Portugal | 36 | 42 | -6 |
| Latvia | 37 | 22 | +15 |
| Germany | 38 | 68 | -30 |
| Bolivia | 39 | 52 | -13 |
| Sweden | 40 | 57 | -17 |
| Uruguay | 41 | 47 | -6 |
| Austria | 42 | 56 | -14 |
| Panama | 43 | 51 | -8 |
| Serbia | 44 | 59 | -15 |
| Trinidad and Tobago | 45 | 37 | +8 |
| Switzerland | 46 | 64 | -18 |
| Turkey | 47 | 84 | -37 |
| The Netherlands | 48 | 71 | -23 |
| Costa Rica | 49 | 61 | -12 |
| Suriname | 50 | 35 | +15 |
| French Polynesia | 51 | 28 | +23 |
| Lebanon | 52 | 66 | -14 |
| Martinique | 53 | 33 | +20 |
| Jordan | 54 | 72 | -18 |
| Gibraltar | 55 | 12 | +43 |
| San Marino | 56 | 13 | +43 |
| Malaysia | 57 | 85 | -28 |
| Guadeloupe | 58 | 39 | +19 |
| Namibia | 59 | 65 | -6 |
| The Bahamas | 60 | 45 | +15 |
| Ireland | 61 | 74 | -13 |
| Estonia | 62 | 60 | +2 |
| Honduras | 63 | 80 | -17 |
| Canada | 64 | 94 | -30 |
| Guatemala | 65 | 89 | -24 |
| Luxembourg | 66 | 62 | +4 |
| Albania | 67 | 76 | -9 |
| Indonesia | 68 | 118 | -50 |
| Guyana | 69 | 69 | 0 |
| Israel | 70 | 87 | -17 |
| Belize | 71 | 58 | +13 |
| Grenada | 72 | 46 | +26 |
| Saint Lucia | 73 | 53 | +20 |
| Andorra | 74 | 44 | +30 |
| Aruba | 75 | 50 | +25 |
| Azerbaijan | 76 | 92 | -16 |
| Botswana | 77 | 81 | -4 |
| Palestine | 78 | 88 | -10 |
| Sri Lanka | 79 | 101 | -22 |
| Eswatini | 80 | 78 | +2 |
| Bermuda | 81 | 49 | +32 |
| Libya | 82 | 93 | -11 |
| India | 83 | 128 | -45 |
| Kazakhstan | 84 | 102 | -18 |
| Cuba | 85 | 99 | -14 |
| Liechtenstein | 86 | 43 | +43 |
| Iraq | 87 | 114 | -27 |
| Sint Maarten | 88 | 48 | +40 |
| Oman | 89 | 97 | -8 |
| Jamaica | 90 | 91 | -1 |
| Philippines | 91 | 121 | -30 |
| Malta | 92 | 79 | +13 |
| Seychelles | 93 | 67 | +26 |
| French Guiana | 94 | 77 | +17 |
| Bahrain | 95 | 96 | -1 |
| Curaçao | 96 | 75 | +21 |
| Saint Martin | 97 | 63 | +34 |
| Belarus | 98 | 113 | -15 |
| Antigua and Barbuda | 99 | 73 | +26 |
| New Caledonia | 100 | 82 | +18 |
| El Salvador | 101 | 112 | -11 |
| Fiji | 102 | 98 | +4 |
| Mongolia | 103 | 108 | -5 |
| Barbados | 104 | 86 | +18 |
| British Virgin Islands | 105 | 70 | +35 |
| Denmark | 106 | 116 | -10 |
| Kuwait | 107 | 115 | -8 |
| Morocco | 108 | 124 | -16 |
| Nepal | 109 | 125 | -16 |
| Myanmar | 110 | 127 | -17 |
| Vietnam | 111 | 131 | -20 |
| Mauritius | 112 | 111 | +1 |
| Thailand | 113 | 132 | -19 |
| Monaco | 114 | 83 | +31 |
| Cabo Verde | 115 | 107 | +8 |
| Kyrgyzstan | 116 | 123 | -7 |
| Dominican Republic | 117 | 126 | -9 |
| Mayotte | 118 | 104 | +14 |
| Cyprus | 119 | 119 | 0 |
| Isle of Man | 120 | 95 | +25 |
| St. Vincent Grenadines | 121 | 100 | +21 |
| Zimbabwe | 122 | 130 | -8 |
| Channel Islands | 123 | 106 | +17 |
| The Caribbean Netherlands | 124 | 90 | +34 |
| Saudi Arabia | 125 | 137 | -12 |
| Réunion | 126 | 122 | +4 |
| Egypt | 127 | 142 | -15 |
| Maldives | 128 | 120 | +8 |
| Dominica | 129 | 109 | +20 |
| Turks and Caicos | 130 | 103 | +27 |
| Bangladesh | 131 | 151 | -20 |
| Finland | 132 | 134 | -2 |
| Lesotho | 133 | 133 | 0 |
| Afghanistan | 134 | 147 | -13 |
| Saint Kitts and Nevis | 135 | 117 | +18 |
| Venezuela | 136 | 145 | -9 |
| Japan | 137 | 154 | -17 |
| Wallis and Futuna | 138 | 105 | +33 |
| UAE | 139 | 141 | -2 |
| Zambia | 140 | 144 | -4 |
| Pakistan | 141 | 158 | -17 |
| Norway | 142 | 138 | +4 |
| St. Barth | 143 | 110 | +33 |
| Cambodia | 144 | 149 | -5 |
| Qatar | 145 | 140 | +5 |
| Algeria | 146 | 156 | -10 |
| Syria | 147 | 153 | -6 |
| Mauritania | 148 | 148 | 0 |
| Sao Tome and Principe | 149 | 136 | +13 |
| Brunei | 150 | 139 | +11 |
| S. Korea | 151 | 164 | -13 |
| Malawi | 152 | 161 | -9 |
| Djibouti | 153 | 146 | +7 |
| Singapore | 154 | 155 | -1 |
| Kenya | 155 | 166 | -11 |
| Senegal | 156 | 162 | -6 |
| Anguilla | 157 | 129 | +28 |
| Comoros | 158 | 150 | +8 |
| Faeroe Islands | 159 | 135 | +24 |
| The Gambia | 160 | 157 | +3 |
| Rwanda | 161 | 165 | -4 |
| Gabon | 162 | 159 | +3 |
| Australia | 163 | 168 | -5 |
| Sudan | 164 | 171 | -7 |
| Uganda | 165 | 172 | -7 |
| Equatorial Guinea | 166 | 160 | +6 |
| Somalia | 167 | 169 | -2 |
| Ethiopia | 168 | 179 | -11 |
| Cameroon | 169 | 173 | -4 |
| Yemen | 170 | 175 | -5 |
| Cayman Islands | 171 | 152 | +19 |
| Mozambique | 172 | 178 | -6 |
| Haiti | 173 | 174 | -1 |
| Timor-Leste | 174 | 167 | +7 |
| Papua New Guinea | 175 | 177 | -2 |
| Iceland | 176 | 163 | +13 |
| Angola | 177 | 181 | -4 |
| Montserrat | 178 | 143 | +35 |
| Congo | 179 | 176 | +3 |
| Guinea-Bissau | 180 | 170 | +10 |
| Uzbekistan | 181 | 182 | -1 |
| Liberia | 182 | 180 | +2 |
| Ghana | 183 | 184 | -1 |
| Madagascar | 184 | 186 | -2 |
| Taiwan | 185 | 185 | 0 |
| Laos | 186 | 183 | +3 |
| Mali | 187 | 188 | -1 |
| Ivory Coast | 188 | 192 | -4 |
| Guinea | 189 | 190 | -1 |
| Nicaragua | 190 | 187 | +3 |
| Togo | 191 | 189 | +2 |
| Hong Kong | 192 | 191 | +1 |
| Nigeria | 193 | 197 | -4 |
| CAR | 194 | 193 | +1 |
| DRC | 195 | 200 | -5 |
| Eritrea | 196 | 194 | +2 |
| Burkina Faso | 197 | 196 | +1 |
| Tanzania | 198 | 202 | -4 |
| Sierra Leone | 199 | 195 | +4 |
| Benin | 200 | 198 | +2 |
| Tajikistan | 201 | 199 | +2 |
| Niger | 202 | 203 | -1 |
| South Sudan | 203 | 201 | +2 |
| Chad | 204 | 204 | 0 |
| New Zealand | 205 | 205 | 0 |
| China | 206 | 207 | -1 |
| Burundi | 207 | 209 | -2 |
| Bhutan | 208 | 206 | +2 |
| Vanuatu | 209 | 208 | +1 |
| Western Sahara | 210 | 210 | 0 |

**Table S3.** A list of countries ranked according to their CO_2_ emissions standardized by the use of linear regression, by *per* *capita* ratios, and the difference between the two approaches.

| **Country Name** | **Ranking based on residuals** | **Ranking based on *per capita* ratios** | **Difference in order** |
| --- | --- | --- | --- |
| Qatar | 1 | 1 | 0 |
| Montenegro | 2 | 5 | -3 |
| Kuwait | 3 | 2 | +1 |
| Trinidad and Tobago | 4 | 4 | 0 |
| United Arab Emirates | 5 | 3 | +2 |
| Oman | 6 | 7 | -1 |
| Canada | 7 | 6 | +1 |
| Brunei | 8 | 12 | -4 |
| Luxembourg | 9 | 14 | -5 |
| Bahrain | 10 | 11 | -1 |
| Australia | 11 | 9 | +2 |
| Estonia | 12 | 13 | -1 |
| Gibraltar | 13 | 19 | -6 |
| Falkland Islands | 14 | 22 | -8 |
| Saudi Arabia | 15 | 10 | +5 |
| United States | 16 | 8 | +8 |
| Turkmenistan | 17 | 15 | +2 |
| Kazakhstan | 18 | 16 | +2 |
| South Korea | 19 | 18 | +1 |
| Iceland | 20 | 26 | -6 |
| Taiwan | 21 | 20 | +1 |
| The Bahamas | 22 | 27 | -5 |
| Russia | 23 | 17 | +6 |
| Czech Republic | 24 | 23 | +1 |
| Bermuda | 25 | 40 | -15 |
| Japan | 26 | 21 | +5 |
| The Netherlands | 27 | 25 | +2 |
| Germany | 28 | 24 | +4 |
| Finland | 29 | 30 | -1 |
| Malaysia | 30 | 29 | +1 |
| Singapore | 31 | 35 | -4 |
| New Caledonia | 32 | 43 | -11 |
| Austria | 33 | 34 | -1 |
| Belgium | 34 | 32 | +2 |
| Ireland | 35 | 37 | -2 |
| Norway | 36 | 36 | 0 |
| Libya | 37 | 39 | -2 |
| Iran | 38 | 31 | +7 |
| Israel | 39 | 38 | +1 |
| Poland | 40 | 33 | +7 |
| Bosnia and Herzegovina | 41 | 42 | -1 |
| China | 42 | 28 | +14 |
| Martinique | 43 | 55 | -12 |
| New Zealand | 44 | 45 | -1 |
| Bulgaria | 45 | 44 | +1 |
| Slovenia | 46 | 48 | -2 |
| South Africa | 47 | 41 | +6 |
| Slovakia | 48 | 47 | +1 |
| Denmark | 49 | 50 | -1 |
| Belarus | 50 | 46 | +4 |
| Hong Kong | 51 | 52 | -1 |
| Cayman Islands | 52 | 65 | -13 |
| Greece | 53 | 51 | +2 |
| Guadeloupe | 54 | 62 | -8 |
| Mongolia | 55 | 57 | -2 |
| Italy | 56 | 49 | +7 |
| Venezuela | 57 | 53 | +4 |
| Cyprus | 58 | 60 | -2 |
| United Kingdom | 59 | 54 | +5 |
| French Guiana | 60 | 72 | -12 |
| Seychelles | 61 | 79 | -18 |
| Spain | 62 | 56 | +6 |
| Barbados | 63 | 73 | -10 |
| Hungary | 64 | 61 | +3 |
| Ukraine | 65 | 59 | +6 |
| Malta | 66 | 76 | -10 |
| France | 67 | 58 | +9 |
| Macao | 68 | 77 | -9 |
| Grenada | 69 | 83 | -14 |
| Portugal | 70 | 66 | +4 |
| Lithuania | 71 | 74 | -3 |
| Switzerland | 72 | 68 | +4 |
| Serbia | 73 | 69 | +4 |
| Antigua and Barbuda | 74 | 88 | -14 |
| Turkey | 75 | 63 | +12 |
| Argentina | 76 | 64 | +12 |
| Croatia | 77 | 75 | +2 |
| Sweden | 78 | 71 | +7 |
| Chile | 79 | 70 | +9 |
| Iraq | 80 | 67 | +13 |
| North Macedonia | 81 | 84 | -3 |
| Guyana | 82 | 87 | -5 |
| Latvia | 83 | 85 | -2 |
| Romania | 84 | 81 | +3 |
| Thailand | 85 | 78 | +7 |
| Saint Kitts & Nevis | 86 | 93 | -7 |
| Algeria | 87 | 80 | +7 |
| Suriname | 88 | 90 | -2 |
| French Polynesia | 89 | 92 | -3 |
| Mexico | 90 | 82 | +8 |
| Uzbekistan | 91 | 86 | +5 |
| Azerbaijan | 92 | 89 | +3 |
| British Virgin Islands | 93 | 106 | -13 |
| Saint Lucia | 94 | 97 | -3 |
| St. Vincent & Grenadines | 95 | 102 | -7 |
| Lebanon | 96 | 91 | +5 |
| Jamaica | 97 | 94 | +3 |
| Belize | 98 | 104 | -6 |
| Botswana | 99 | 95 | +4 |
| Panama | 100 | 96 | +4 |
| Gabon | 101 | 103 | -2 |
| Aruba | 102 | 116 | -14 |
| Cuba | 103 | 98 | +5 |
| Dominica | 104 | 118 | -14 |
| Tunisia | 105 | 100 | +5 |
| Maldives | 106 | 115 | -9 |
| Mauritius | 107 | 112 | -5 |
| Tonga | 108 | 121 | -13 |
| Ecuador | 109 | 105 | +4 |
| Jordan | 110 | 111 | -1 |
| Palau | 111 | 129 | -18 |
| Egypt | 112 | 101 | +11 |
| North Korea | 113 | 109 | +4 |
| Bhutan | 114 | 119 | -5 |
| Dominican Republic | 115 | 113 | +2 |
| Brazil | 116 | 99 | +17 |
| Vietnam | 117 | 108 | +9 |
| Saint Helena | 118 | 134 | -16 |
| Syria | 119 | 114 | +5 |
| Georgia | 120 | 117 | +3 |
| Cook Islands | 121 | 133 | -12 |
| Anguilla | 122 | 135 | -13 |
| Indonesia | 123 | 110 | +13 |
| Moldova | 124 | 122 | +2 |
| Fiji | 125 | 127 | -2 |
| India | 126 | 107 | +19 |
| Uruguay | 127 | 123 | +4 |
| Peru | 128 | 120 | +8 |
| Albania | 129 | 128 | +1 |
| Turks and Caicos | 130 | 139 | -9 |
| Equatorial Guinea | 131 | 131 | 0 |
| Bolivia | 132 | 124 | +8 |
| Costa Rica | 133 | 130 | +3 |
| Namibia | 134 | 132 | +2 |
| Morocco | 135 | 125 | +10 |
| Djibouti | 136 | 137 | -1 |
| Colombia | 137 | 126 | +11 |
| Armenia | 138 | 136 | +2 |
| Saint Pierre & Miquelon | 139 | 148 | -9 |
| Réunion | 140 | 141 | -1 |
| Philippines | 141 | 138 | +3 |
| Kyrgyzstan | 142 | 143 | -1 |
| Guatemala | 143 | 140 | +3 |
| Papua New Guinea | 144 | 144 | 0 |
| El Salvador | 145 | 145 | 0 |
| Angola | 146 | 142 | +4 |
| Congo | 147 | 147 | 0 |
| Honduras | 148 | 149 | -1 |
| Yemen | 149 | 150 | -1 |
| Paraguay | 150 | 152 | -2 |
| Sri Lanka | 151 | 151 | 0 |
| Pakistan | 152 | 146 | +6 |
| Samoa | 153 | 155 | -2 |
| Nicaragua | 154 | 153 | +1 |
| Zimbabwe | 155 | 154 | +1 |
| Tajikistan | 156 | 156 | 0 |
| Laos | 157 | 157 | 0 |
| Mauritania | 158 | 159 | -1 |
| Benin | 159 | 158 | +1 |
| Eswatini | 160 | 163 | -3 |
| Senegal | 161 | 160 | +1 |
| Solomon Islands | 162 | 165 | -3 |
| Ghana | 163 | 162 | +1 |
| Vanuatu | 164 | 169 | -5 |
| Bangladesh | 165 | 161 | +4 |
| Kiribati | 166 | 171 | -5 |
| Nigeria | 167 | 164 | +3 |
| Côte d'Ivoire | 168 | 166 | +2 |
| Cambodia | 169 | 167 | +2 |
| Timor-Leste | 170 | 172 | -2 |
| Cameroon | 171 | 168 | +3 |
| Western Sahara | 172 | 176 | -4 |
| South Sudan | 173 | 170 | +3 |
| Kenya | 174 | 173 | +1 |
| Sudan | 175 | 174 | +1 |
| Myanmar | 176 | 175 | +1 |
| Togo | 177 | 177 | 0 |
| Nepal | 178 | 178 | 0 |
| Afghanistan | 179 | 179 | 0 |
| Haiti | 180 | 180 | 0 |
| Sao Tome & Principe | 181 | 182 | -1 |
| Zambia | 182 | 181 | +1 |
| Puerto Rico | 183 | 184 | -1 |
| Mozambique | 184 | 183 | +1 |
| Eritrea | 185 | 186 | -1 |
| Cabo Verde | 186 | 191 | -5 |
| Tanzania | 187 | 185 | +2 |
| Guinea | 188 | 187 | +1 |
| Liberia | 189 | 188 | +1 |
| Guinea-Bissau | 190 | 190 | 0 |
| Sierra Leone | 191 | 189 | +2 |
| Lesotho | 192 | 192 | 0 |
| Comoros | 193 | 197 | -4 |
| Uganda | 194 | 193 | +1 |
| Burkina Faso | 195 | 194 | +1 |
| Madagascar | 196 | 195 | +1 |
| Rwanda | 197 | 196 | +1 |
| Central African Republic | 198 | 199 | -1 |
| The Gambia | 199 | 201 | -2 |
| Malawi | 200 | 202 | -2 |
| Chad | 201 | 198 | +3 |
| Ethiopia | 202 | 200 | +2 |
| Niger | 203 | 204 | -1 |
| Burundi | 204 | 203 | +1 |
| Mali | 205 | 206 | -1 |
| Somalia | 206 | 207 | -1 |
| DR Congo | 207 | 205 | +2 |
| Faeroe Islands | 208 | 208 | 0 |
| Greenland | 209 | 209 | 0 |
